# Assessing Inter-Individual Genetic Variability in Peach Sugar Metabolism through Reliable Parameter Estimation of a Kinetic Model

**DOI:** 10.1101/2023.08.10.552764

**Authors:** Hussein Kanso, Mohamed-Mahmoud Memah, Valentina Baldazzi, Bénédicte Quilot-Turion, Charlotte Baey

## Abstract

Integrating genetic information into dynamical models is key to understand variations among genotypes and improve crop performances face to current agronomic and ecological constraints. A first and necessary step in building gene-to-phenotype models is the calibration of a large number of genotypes. We compared here two different strategies for the calibration of an Ordinary Differential Equations (ODE) kinetic model simulating the accumulation of different sugars during peach fruit development. First, the model was calibrated for each genotype independently using a Genotype-Based (GB) strategy. Two formulations of the problem have been tested, either as a Single-Objective Optimization (GBS) problem or as a Multi-Objective Optimization (GBM) problem. Second, the model was calibrated for all genotypes simultaneously using a Population-Based (PB) strategy. The two strategies were first applied to a set of simulated data and then to a real dataset derived from an interspecific population of 106 peach genotypes. Results showed that the GB strategy allowed for a high goodness of fit for most genotypes, especially in the GBS formulation. However, the estimated parameters suffered from a lack of practical identifiability as independent repetitions of the estimation algorithm did not always converge to the same value for most genotypes. The PB calibration strategy overcame this issue showing a good identifiability of the population parameter values, a goodness of fit comparable to the one obtained with the GB strategy and a good characterisation of parameter variations within the progeny, which is a key to assess the inter-individual genetic variability. These results are an important step towards the development of reliable gene-to-phenotype models.

## 1 Introduction

Predicting genotype-to-phenotype relationships under contrasting environments is a big challenge for plant biology and breeding. Process-based ecophysiological models are considered as powerful tools to deal with such a challenge (Martre et al., 2015) as they are able to decompose plant phenotypic traits into developmental, architectural and environmental factors. Within this framework, differences among genotypic responses are usually encoded into a set of parameters values, considered as the fingerprint of the underlying genetic information (Martre et al., 2015). The aim of gene-to-phenotype modelling is to make this link explicit, by expressing the value of genetic parameters as a function of specific genetic loci (QTL) or allelic combinations of genes, depending on the available information. Ultimately, gene-to-phenotype models can be used to explore trait correlations in a population and to analyse Genotype (G) x Environment (E) x Management (M) interactions, in the perspective of virtual breeding.

The construction of gene-to-phenotype models is challenging as it relies on the calibration of a large number of genotypes, usually with few data available. Moreover, process-based models are generally nonlinear and involve a large number of parameters among which correlations may exist (Chou and Voit, 2009; Gutenkunst et al., 2007). In such cases, non-identifiability issues may arise, when parameter values can not be unambiguously inferred from available data and multiple parameter sets provide equally good fits of the data. Although in many biological applications, useful insights may be derived from model predictions rather than parameter values, uncertainty in parameter estimates poses a problem for genetic analysis, in which the parameters values are expected to be the genetic fingerprint of the system (Hass et al., 2017; Becker et al., 2010). Moreover, spurious correlations among parameter estimates along with sloppy directions, may prevent the identification of true genetic correlations due to epistastic effects or common regulatory mechanisms. In general, a reliable estimate of the parameters depends on the structure of the model, the quality of the observations and the choice of the calibration strategy. Although model reduction approaches can be effectively used to improve model identifiability (Maiwald et al., 2016; Chou and Voit, 2009; Snowden et al., 2017), we focus here on the last point, namely the formalization and implementation of the calibration process.

In this work, we compared two different strategies for the calibration of an ODE kinetic model simulating the accumulation of different sugars during peach fruit development (Desnoues et al., 2018; Kanso et al., 2020). The first strategy corresponded to the common approach in plant modeling community: parameter estimation was addressed for each genotype independently (Genotype-based, GB), and the obtained estimates were compared in order to identify the genotypic parameters of the system (Bertin et al., 2010). Examples of this approach can be found in the study of many plant processes, including leaf elongation (Reymond et al., 2003), plant development (Yin et al., 2000, 2005; Messina et al., 2006), phenology (Nakagawa et al., 2005; Ravi Kumar et al., 2009), nitrogen adaptation (Laperche et al., 2006), fruit growth (Constantinescu et al., 2016) and quality (Quilot et al., 2005; Prudent et al., 2011). This strategy has been largely employed as it has the advantage to be straight forward: it can be performed with any kind of optimization algorithm, applied to any model and can be easily extended to newly acquired genotypes without recalibrating all the dataset. However, as estimations are performed for each genotype independently without sharing any information, the estimated parameters using GB approach could show large variations for the whole population. In the second strategy, we calibrated the model using a Population-based (PB) approach based on mixed-effects models, which assumes that individual parameters, defined for each genotype, are independent and identically distributed random variables following a common probability distribution (Davidian and Giltinan, 2003; Baey et al., 2016). Accordingly, the genetic population is treated as a whole and the model is calibrated for all genotypes simultaneously. The target of the estimation shifts from the individual profile to the average behaviour, in terms of parameter distribution across the population. Individual parameters values can be empirically obtained as random variation around the estimated mean population curve. PB approach has been largely used in phamacokinetics to characterize the inter-individual variability of observed drug-response dynamics (Donnet and Samson, 2013; Owen and Fiedler-Kelly, 2014). It has been recently applied to the calibration and improvement of functional-structural plant growth models, in relation to plant genetic diversity (Perez et al., 2016; Baey et al., 2016, 2018).

Although both strategies have proven successful in a number of cases, to our knowledge, they have never been compared. The aim of this paper is to test their capacities to assess the interindividual variability of parameter values in the context of genetic studies, using the model of sugar metabolism in peach fruit as a case study. To this aim, the two strategies were first applied on a set of simulated data to evaluate the impact of each calibration strategy on the estimation of individual genotypic parameters, in terms of accuracy and uniqueness of the solution. Then we applied each approach to a real dataset derived from an interspecific population of 106 peach genotypes. For the GB strategy, two formulations of the calibration problem were evaluated, either as a Single-Objective Optimization (GBS) problem or as a Multi-Objective Optimization (GBM). In the first case i.e. GBS formulation, the estimation process results in a unique value for the objective function. Hopefully, the value of this objective function corresponds to a unique set of parameter values for each genotype of the progeny. On the other hand, using the GBM formulation, an entire set of trade-offs between antagonist objective functions (Pareto front) is identified for each genotype, among which the desired parameter set has to be selected based on some other criteria. This choice could be done using graphical tools, decision-making methods, or some particular heuristics designed for a particular classification problem (Coello et al., 2007; Akdemir et al., 2019). This allows for a deeper analysis of possible trade-offs among physiological traits, in the case of complex phenotypes.

The paper is organized as follows: Section 2 details the mathematical model, Section 3 introduces the problem formulation and the optimisation methods, Section 4 presents the simulated and experimental datasets used for the study and Section 5 describes the indicators used for assessing the quality of the two calibration strategies. Results are reported and discussed in Section 6 from the point of view of gene-to-phenotype modeling. Finally, a discussion concludes the paper in Section 7.

## 2 Mathematical model

The model used in this paper is the one proposed by Kanso et al. (2020) as a reduced version of the model previously developed by Desnoues et al. (2018). The model describes the accumulation of four different sugars (sucrose, glucose, fructose, and sorbitol) in peach fruit during its development, as a system of ordinary differential equations. Carbon enters the fruit from the plant sap which is transformed by a metabolic network, including enzymatic reactions and transport mechanisms between the cytosol and the vacuole (see Figure 1). A total of 17 reactions are included in the model and represented by linear flows. The model makes explicit use of experimental data to describe the temporal evolution of fruit carbon supply and fruit volume. To this aim, measured fruit dry and fresh masses are represented by genotype-specific temporal functions and provided as input to the model (see Supplementary section S1). The model contains 30 parameters, among which 21 are considered as known (measured or taken from literature) and identical for all genotypes. This work focuses on the estimation of the remaining 9 parameters (Table 1), supposed genotype-dependent.

**Table 1:** Description of the 9 unknown genotype-dependent parameters of the sugar model.

| Parameter | Description | Range | Unit |
| --- | --- | --- | --- |
| $LHx$ | sucrose proportion hydrolyzed in the apoplasm | $]0, 1]$ | |
| $RAI$ | coefficient of the transfer function between sucrose and (glucose +fructose) under action of acid invertase (AI) enzyme | $]0, 0.1]$ | $day^{-1}$ |
| $KFk$ | fructokinase (FK) affinity | $]0, 100]$ | $mggFW^{-1}$ |
| $KHk$ | hexokinase (HK) affinity | $]0-500]$ | $mggFW^{-1}$ |
| $OCp$ | coefficient of the transfer function between hexoses phosphate and other compounds | $[250, 1500]$ | $day^{-1}$ |
| $RSS$ | coefficient of the transfer function between hexoses phosphate and sucrose | $]0, 1000]$ | $day^{-1}$ |
| $TAS$ | coefficient of sucrose transport (active import) from cytosol to vacuole | $]0, 400]$ | $mggFW^{-1}day^{-1}$ |
| $TPF$ | coefficient of fructose passive transport between cytosol and vacuole and in the opposite direction | $]0, 150]$ | $mggFW^{-1}day^{-1}$ |
| $RSO$ | coefficient of the transfer function between sorbitol and glucose under action of sorbitol oxydase (SO) enzyme | $]0, 10]$ | $day^{-1}$ |

**Figure 1:**
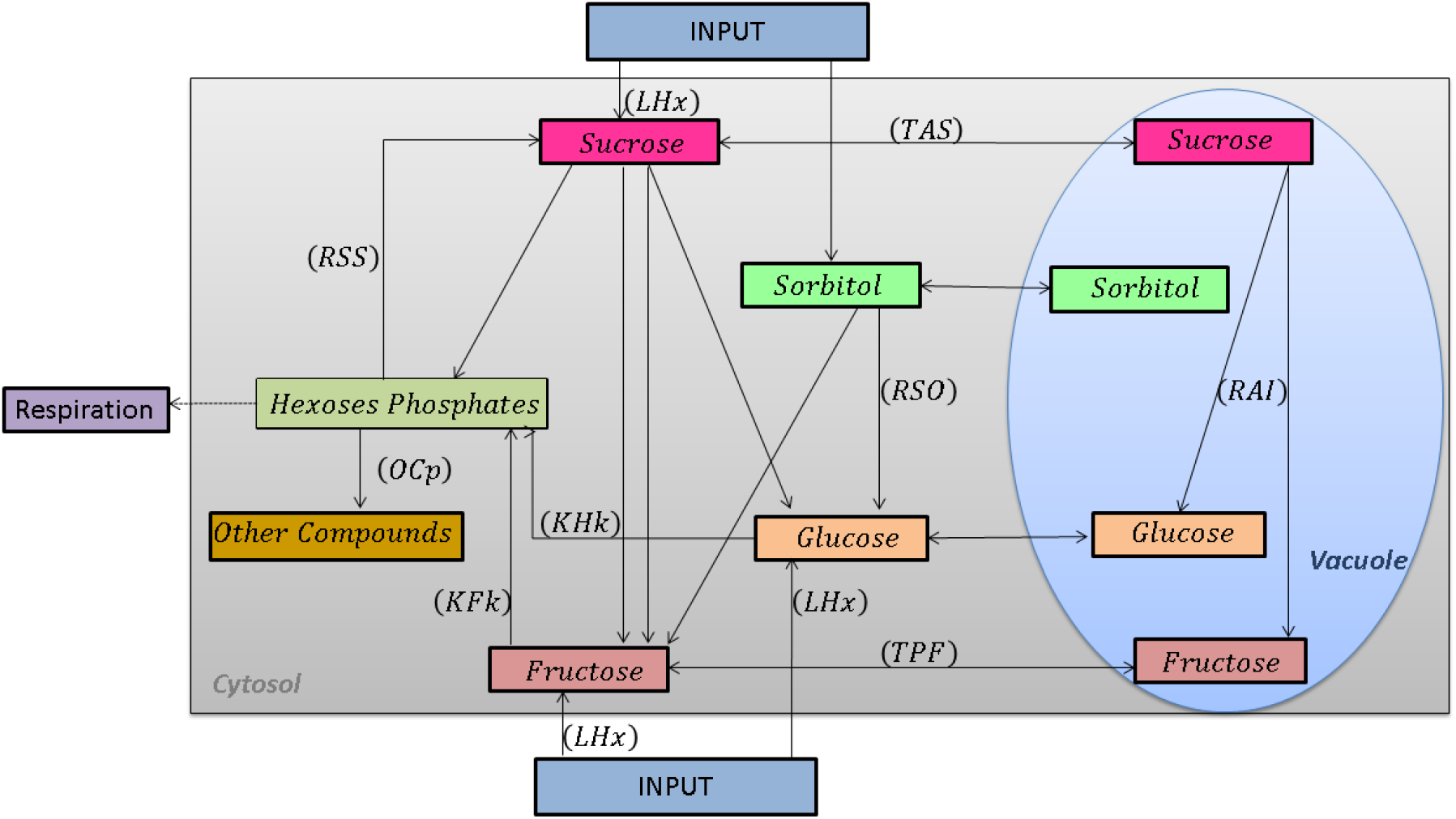
Schematic network of the reduced model of sugar accumulation in the peach fruit. Arrows represent carbon flows. The corresponding kinetic equations are presented in Kanso et al. (2020).

## 3 Problem formulations and calibration strategies

The aim of this work was to estimate the value of the 9 genotype-dependent parameters using data on sugar concentrations, at six developmental stages, from a large number of genotypes. Two calibration strategies were compared i) parameters were estimated independently for each genotype (GB strategy) and ii) parameters were simultaneously estimated for the whole progeny (PB strategy). Figure 2 gives a schematic representation of the calibration steps, detailed below. We consider a common framework for the two calibration strategies. Let us denote by *y*_*ijk*_, *i* = 1, …, *N*_*jk*_, *j* = 1, …, 4, *k* = 1, …, *G* the *i*-th observation of the *j*-th sugar concentration for genotype *k*, and by ℳ_*ij*_ the corresponding model prediction.*N*_*jk*_ is the number of measurements for genotype *k* and sugar *j*,

**Figure 2:**
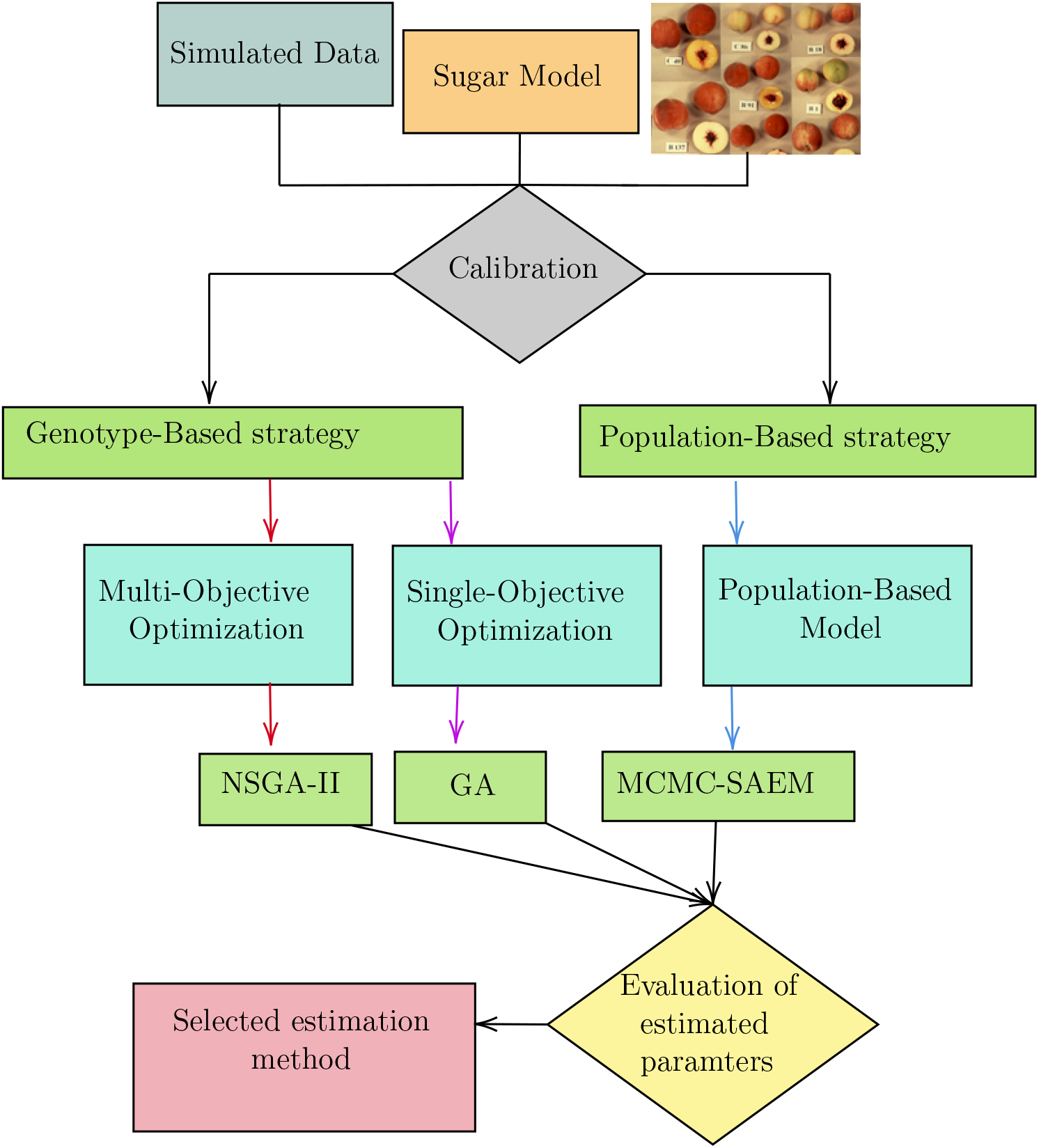
Graphical representation of the proposed steps to calibrate the sugar model. Two strategies were used to calibrate the model: 1) Genotype-Based strategy where two formulations were considered: Single-Objective Optimisation (GBS) and Multi-Objective Optimisation (GBM), 2) Population-Based (PB) strategy. To estimate the unknown 9 genotype-dependent parameters (See Table 1), 3 algorithms were used : GA for GBS, NSGA-II for GBM and MCMC-SAEM for PB.

We then denote by 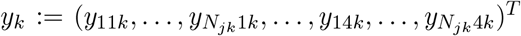 the vector of all observations for genotype *k*, and by ℳ_*k*_ the associated vector of all model predictions for genotype *k*. More specifically, observations are gathered per sugar type. We assume the following model:

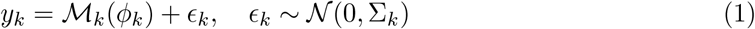

where *ϕ*_*k*_ is the vector of genotypic parameters to be estimated, *ϵ*_*k*_ is the residual error assumed to follow a Gaussian distribution with mean 0 and covariance matrix Σ_*k*_. Three parsimonious models were driven from the above one based on some hypotheses detailed in the following subsections.

### 3.1 Model’s calibration using genotype-based strategy

In the framework of this first strategy, we used two alternative formulations of the model calibration as a single-objective optimization (GBS), and as a multi-objective optimization (GBM) problem.

#### Calibration as Single-Objective Optimisation (GBS)

The single-objective optimisation formulation combines observations of the four sugars together into a single objective defined as the sum of the squared errors of each sugar. Using equation (1), we assume that 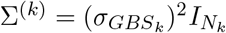, where 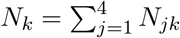 is the total number of measurements for genotype *k* and *I*_*p*_ is the identity matrix of size *p × p*, i.e. we assume that the variance of the error term is independent of the sugar type. We denote by *θ*_*GBS*,*k*_ = (*ϕ*_*GBS*,*k*_, *σ*_*GBS*,*k*_) the vector of parameters for genotype *k*. The log-likelihood function for genotype *k* is denoted by *ℓ*_*GBS*,*k*_ and assuming independence between data within a given genotype, is defined as follows:

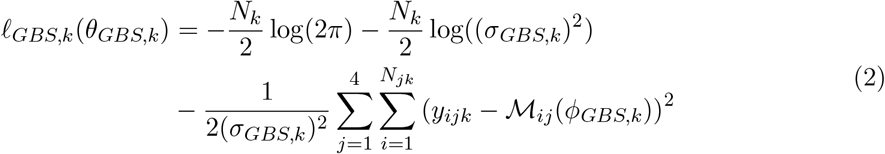

A maximum log-likelihood estimator of *θ*_*GBS*,*k*_ is defined as a solution to the maximization problem:

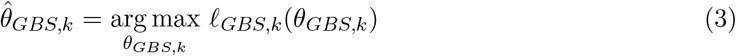

In this Gaussian case, the maximum log-likelihood estimator is equivalent to the ordinary least-square estimator:

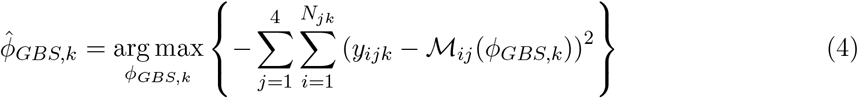

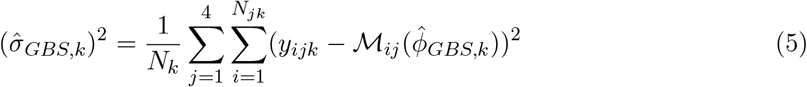

#### Calibration as Multi-Objective Optimisation (GBM)

In the multi-objective optimisation formulation, the optimisation is performed on a vector of objective functions. Contrary to GBS, each sugar is considered independently. Thus, the problem can be decomposed into four objective functions, each of which addressing one sugar. These functions are considered as separate objectives to be optimised independently. The idea is to estimate the parameters by finding a compromised solution between objective functions. In this context, and using equation (1), we have Σ_*k*_ = diag(Σ_1*k*_, …, Σ_4*k*_), where 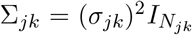, i.e. we assume that the variance of the error term depends on the sugar type.

We denote by *θ*_*GBM*,*jk*_ = (*ϕ*_*GBM*,*k*_, *σ*_*GBM*,*jk*_) the vector of parameters for genotype *k* and sugar *j* and by *θ*_*GBM*,*k*_ = (*ϕ*_*GBM*,*k*_, (*σ*_*GBM*,*jk*_)_*j*=1,…,4_) the vector of parameters for genotype *k*. The log-likelihood function for genotype *k* and sugar *j* is denoted by *ℓ*_*GBM*,*j*,*k*_, and under the assumption of independence between the observations, it is given by:

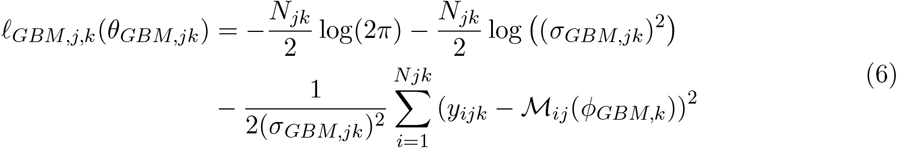

We denote by *ℓ*_*GBM*,*k*_ (*θ*_*GBM*,*k*_) = (*ℓ*_*GBM*,1,*k*_ (*θ*_*GBM*,1,*k*_), …, *ℓ*_*GBM*,4,*k*_ (*θ*_*GBM*,4,*k*_))^*T*^ the vector of log-likelihoods for genotype *k*, and the maximum likelihood estimator for genotype *k* is given by:

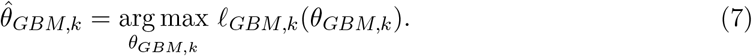

Maximising each log-likelihood *ℓ*_*GBM*,*j*,*k*_ lead to the same explicit formulation for *ϕ*_*GBM*,*k*_ and *σ*_*GBM*,*jk*_ as in equations (4) and (5). Note that here *σ*_*GBM*,*jk*_ depends on *j* as we assumed that the error term depends on the sugar type.

### 3.2 Model calibration using population-based strategy

We used the statistical approach developed by Baey et al. (2018) for a simultaneous parameter estimation of all the genotypes in the population. In the PB strategy, the statistical model is defined as a two-stage hierarchical one. The first stage deals with the intra-individual variability. The observations of each genotype are modeled as in equation (1), with 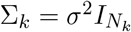.

The second stage deals with the inter-individual variability. More precisely, the individual parameters *ϕ*_*k*_ are considered as random variables. In our case, log transformations were applied to these parameters as they are all strictly positive by definition:

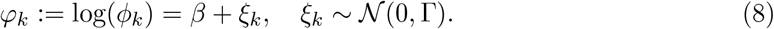

In the sequel, the term ‘random effects’ will refer equivalently to *ϕ*_*k*_ or *φ*_*k*_.

We denote by *θ*_*PB*_ = (*β*, Γ, *σ*_*PB*_) the vector of parameters for the PB model. We assume that the random effects *ϕ*_*k*_ are mutually independent, and that the within-genotype observations are conditionally independent, given the random effects. Thus; the log-likelihood of the PB model is defined as:

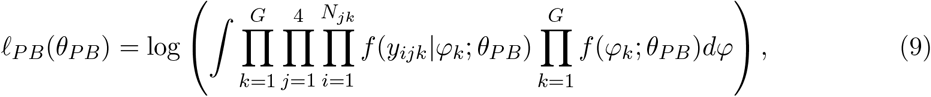

where *f* (*y*_*ijk*_|*φ*_*k*_; *θ*_*PB*_) is the conditional density of the observations *y*_*ijk*_ given the individual parameters, and *f* (*φ*_*k*_; *θ*_*PB*_) is the marginal density of *φ*_*k*_.

Due to the non-linearity of the mathematical model ℳ in equation (1), an analytical computation of the integral (9) is impossible in practice. However, mixed models can be seen as a special case of incomplete data models, with (*y*_*ijk*_) being the observed data, and the random effects *φ*_*k*_ being the missing data. In this case, maximum likelihood estimation can be done using an EM-type algorithm (Dempster et al., 1977).

Confidence intervals can be obtained for each component of *θ*_*PB*_, using a parametric Bootstrap approach once the EM-type algorithm has converged (Efron and Tibshirani, 1993).

### 3.3 Optimisation algorithms and their settings

For the GB strategy, we used two genetic algorithms for the numerical optimisation of the objective functions: GA (Goldberg, 1989; Conn et al., 1991, 1997) for the GBS formulation and NSGA-II (Deb et al., 2002) for GBM formulation.

For the PB strategy, we used a MCMC-SAEM algorithm (Kuhn and Lavielle, 2004, 2005), coupling a Markov chain Monte Carlo algorithm (Hastings, 1970) with a Stochastic Approximation Expectation-Maximization (Robbins and Monro, 1951). This method allows for an estimation of *θ*_*PB*_, but individual estimates for *ϕ*_*k*_ (or equivalently *φ*_*k*_) can also be obtained as a by-product of the algorithm. More details on the implementation of this algorithm in the context of plant growth modelling can be found in Baey et al. (2018).

Given the stochastic nature of algorithms, the estimation procedure was repeated multiple times to ensure the good exploration of the parameters space and test the robustness of the approach. For GA and NSGA-II, the estimation procedure was repeated 10 (real data) to 20 (simulated data) times for each genotype *k*, whereas for MCMC-SAEM, the estimation procedure was repeated 5 (simulated data) to 10 (real data) times, for all genotypes simultaneously. In the case of GBS, the results of all repetitions were combined together. Then, the solution having the maximum likelihood was selected as the reference estimation, for each genotype *k*. In addition, the solutions having likelihoods highest than 95th percentile of all likelihood values were kept to analyse the intra-individual variability of the estimated parameter values. Unlike in GBS, GBM does not yield a single optimal solution but identifies a Pareto-optimal set i.e. solutions corresponding to the best reached trade-offs between our four likelihoods i.e. Pareto front. The issue is then how to select a single reference solution among all these options (Constantinescu et al., 2016). Many methods can be used for this purpose including graphical tools and multicriteria decision making methods. Here, all solutions resulting from the independent repetitions of the calibration process were first pooled together and then filtered in order to identify the final Pareto-optimal set. We then summed the four individual likelihoods for each solution in this Pareto-set and choose solution(s) with highest value of this sum.

In the case of PB, the estimation having the highest likelihood value was selected as the reference solution and used for the subsequent analyses.

## 4 Experimental and simulated data

### 4.1 Simulation study

The proposed calibration strategies were first applied to a set of 𝒮 simulated datasets (𝒮 =100), in order to check the quality and stability of the corresponding estimator. Given the intrinsic difference between the GB and PB strategies, distinct procedures were used to built the corresponding datasets. These procedures are detailed in the following sections.

#### Datasets for the genotype-based strategy

To test the quality of the GB strategy (GBS and GBM formulations), we used the model to build simulated datasets corresponding to repeated noisy measurements of the *same* genotype.

Accordingly, a single set of parameters (see Table 2) and the same model input (see section S1 of the Supplementary Text) were used to simulate the dynamics of the four sugars by means of our ODE model. In order to be close to the experimental limitations, only 6 dates were selected along the dynamics and used as observations for the subsequent calibration. In order to obtain 𝒮 different measurements, a random observation noise with variance *σ* was added to the simulated values, at the considered time points.

**Table 2:** Parameters used to simulate the dynamics of sugar concentrations, in the GB strategy.

|  |  |
| --- | --- |
| LHx | 0.03 |
| RAI | 0.01 |
| KFk | 12.18 |
| KHk | 20.08 |
| OCp | 665.14 |
| RSS | 33.11 |
| TAS | 20.08 |
| TPF | 12.18 |
| RSO | 1.01 |
| $\sigma_k^2$ | 0.05 |

#### Datasets for the population-based strategy

For the population-based approach, 𝒮 populations were generated, each one including *G* = 100 different genotypes. All the nine effects were considered as random and Γ was assumed to be diagonal. The population mean *β* was defined as the logarithm of the values in Table 2 and the variances were equal to 10% of the mean, i.e. diag(Γ) = 0.10 *β*.

For each simulated population, a set of *G* individual parameters was generated according to Eq. 8. Observations were then obtained by running the model according to Eq. 1, with *σ*^2^ = 0.05. As in the previous case, six dates were selected, evenly distributed along the fruit developmental period in order to be close to experimental conditions. For each genotype, model inputs (fruit dry and fresh weights) were randomly assigned as described in section S1.

### 4.2 Experimental data

A progeny of 106 genotypes was used in this study. As described by Quilot et al. (2004), these genotypes come from an inter-specific progeny obtained by two subsequent back-crosses between *Prunus davidiana* (Carr.) P1908 and *Prunus persica* (L.) Batsch ‘Summergrand’ and then ‘Zephyr’. The concentrations of different metabolites, namely sucrose, glucose, fructose, sorbitol, and hexoses phosphates, the fruit flesh fresh mass and the flesh dry matter content, as well as enzymatic capacities (maximal activity) of twelve enzymes were measured at six time points during fruit development, for all genotypes, as described in Desnoues et al. (2014). At each time point, available data consists in 3 biological measurements for 10 genotypes and 1 biological measurement for the remaining 96 genotypes. Genotypes in the studied progeny can be sorted into two classes: those having a ‘standard phenotype’, namely a balanced fructose-to-glucose ratio at maturity, and those having a ‘low fructose phenotype’ due to the lower proportion of fructose compared with glucose at maturity (Desnoues et al., 2018).

## 5 Quality indicators

The different estimation methods were evaluated using several criteria dealing with i) the quality of phenotypic predictions, and ii) the accuracy and variability of the parameter estimates.

### 5.1 Quality of phenotypic predictions

#### Correlation between prediction and observation

Pearson’s linear correlation coefficient Corr_*j*_ was used to quantify the relationship between predictions and observations over the whole peach population, for each sugar *j*:

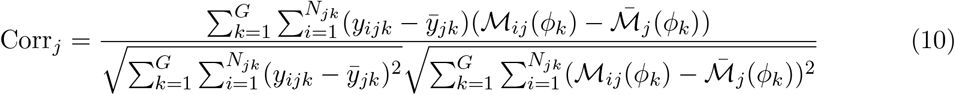

where 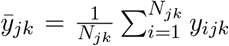 and 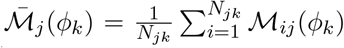. Values of the correlation coefficient can range from –1 to +1. A value of –1 indicates perfect negative correlation, while a value of +1 indicates perfect positive correlation. A value of 0 indicates no correlation between the variables.

#### NRMSE: Normalized Root Mean Square Error

The Normalized Root Mean Square Error (NRMSE) provides a measure of the agreement between model predictions and data all along the dynamics of sugar accumulation. The NRMSE for genotype *k* and sugar *j* was defined as:

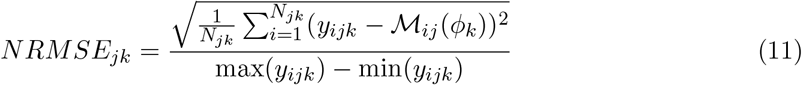

In order to get a comprehensive view of the quality of model predictions, a weighted NRMSE (*aNRMSE*) was constructed at the level of the whole sugar phenotype as

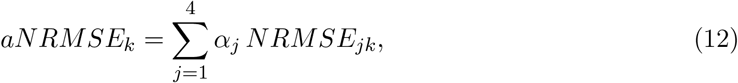

where *NRMSE*_*jk*_ is the Normalized Root Mean Square Error computed for sugar *j* and *α*_*i*_ the relative proportion of sugar *i* in the fruit. *α* = (0.72, 0.13, 0.09, 0.05) for sucrose, glucose, fructose, and sorbitol, respectively.

### 5.2 Quality of parameter estimates

#### Root mean squared Error

The root mean squared error (RMSE) of an estimator 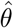 is a measure of its accuracy. It quantifies the risk of using this estimator as a proxy for the real parameter value *θ*, in the framework of parameter estimation. The lower the RMSE value, the more the estimator is reliable.

The RMSE is defined as:

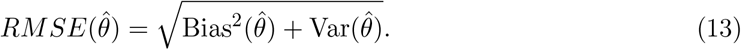

In practice, the bias is estimated as the difference between the average value of our reference estimates (as a proxy of the expected value) and the correct value which we are trying to predict:

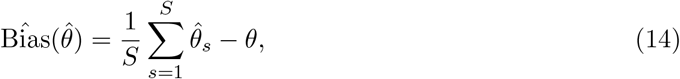

where *ϕ* is the value used to generate the simulated data. The variance is estimated by

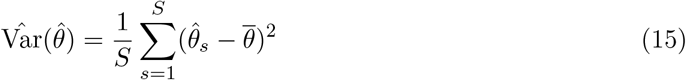

where 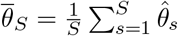 is the mean value of our reference estimates.

The RMSE can be computed for the parameters associated with the virtual genotype generated for the GB strategy (the nine model parameters and the observation noises), and for the parameters associated with the virtual population generated for the PB strategy (i.e. the population parameters but also the *G* individual parameters).

#### Expected Error on individual parameters

In the context of our simulation study, the quality of individual parameters estimation can be assessed by computing the relative distance between the true value *ϕ*_*r*,*k*_ and the estimated value 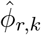 for the *r*-th parameter and the *k*-th genotype. Accordingly, the Expected Error *E*_*r*,*k*_ (in %) was defined as:

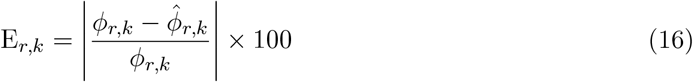

Overall, for each of the 9 parameters, our simulation study yielded *S* values of E_*r*,*k*_ in the case of GB approach and *S × G* values for the PB approach.

#### Relative standard deviation

The relative standard deviation (RSD) was used to quantify the estimate variability of the *r*-th parameter of genotype *k*, over *R* repetitions of the calibration algorithm. RSD is defined as:

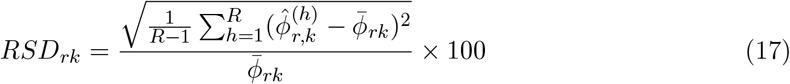

where 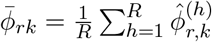 is the average estimate of the considered parameter.

## 6 Results

### 6.1 The Genotype- and Population-based strategies provide comparable fit quality on real data

GB and PB strategies were applied to a population of 106 peach genotypes for which data on fruit sugar composition were available (section 4.2). Figure 3 shows the predictions-observations plots obtained for the four sugars using GB and PB calibration approaches. In general, both estimation strategies provided acceptable predictions for the four sugars, with particular emphasis on sucrose, the dominant sugar in peach fruit. Predictions were less accurate for sorbitol compared to the other sugars, regardless of the calibration method. This can be partially explained by the quality of experimental measurements, which can suffer from a significant error due to the low concentration of sorbitol in the fruit.

**Figure 3:**
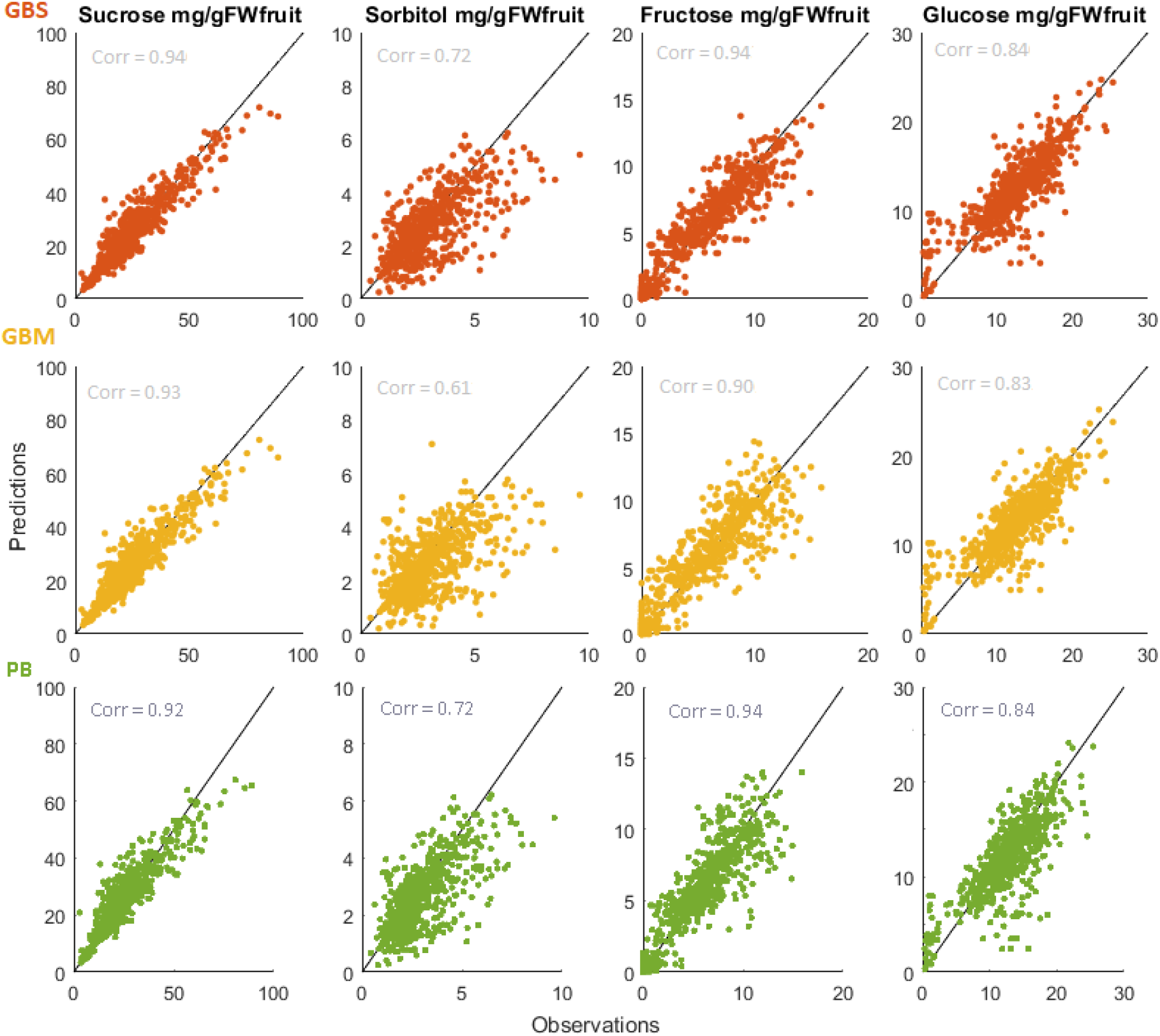
Predictions vs observations using the individual estimates over the peach population. Top: results from (GBS/GA). Middle: results from (GBM/NSGA-II). Bottom: results from (PB/MCMC-SAEM). Corr: Correlations computed between predictions and observations for each sugar and each calibration formulation/algorithm.

Results were confirmed by computing correlation values between predictions and observations for each problem. GBS and PB methods gave similar if not identical results for the prediction of sugar concentrations. GBM performed well on sucrose but was clearly outdistanced on minor sugars. Despite the multi-objective definition of the corresponding likelihood function, the fitting quality of GBM appeared to be driven by the dominant sugar. Moreover, differences in the fitting quality can be observed among individual genotypes. When looking at the prediction of the sugar accumulation dynamics, most aggregate NRSME values range between 0.15 and 0.25, for both GB and PB methods, but outliers exist with large prediction errors, above 0.4 (Fig. 4). In particular, a handful of data points at low glucose concentration were consistently overestimated (Fig. 3), independently of the calibration algorithm. These points suggest the presence of atypical genotypes in the population, for which the underlying model is not well-suited.

**Figure 4:**
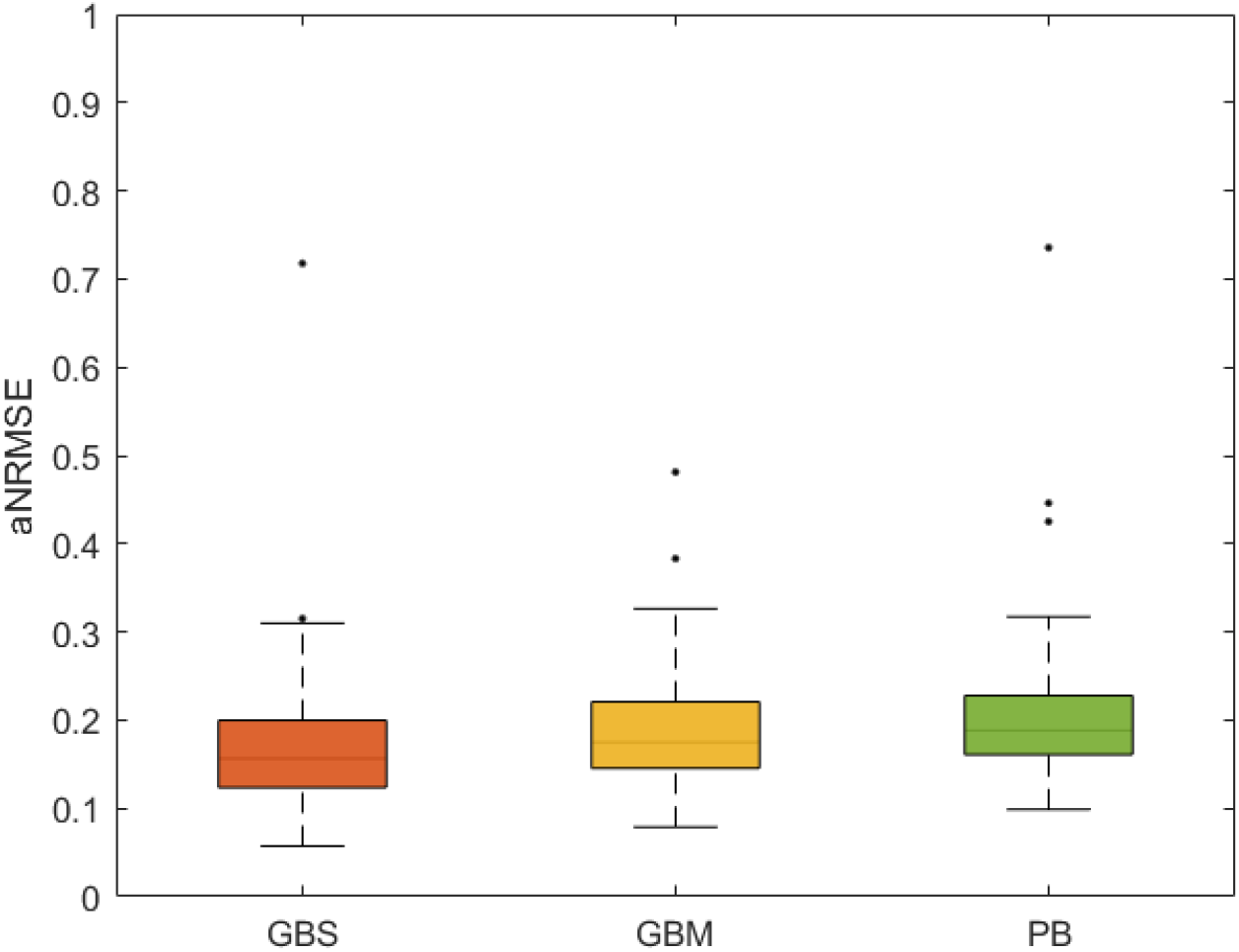
Aggregate Normalized Root Mean Square Errors (aNRMSE) between model predictions and experimental data using GB and PB calibration approaches. The boxplots show the variability of aNRMSE over the peach population.

### 6.2 Population-based strategy provides higher statistical accuracy

In the previous section, we were concerned by the overall goodness-of-fit of each calibration strategy. We now investigate the link between the phenotype and the values of the genetic parameters. Indeed, in order to be used for genetic studies, parameter estimations should be robust across repetitions, providing a unique mapping between the individual phenotype (sugar concentrations) and the underlying parameter values. In this section, we use data from our simulation study to evaluate the performance of each algorithm to estimate the model parameters, based on root mean squared error (RMSE) of the associated statistical estimator. Results are reported in Tables 3.

**Table 3:** Comparison of parameters estimates: means and root mean square error based on 100 simulated datasets.

| Genotype-based |  |  |  |  |  | Population-based |  |  |  |
| --- | --- | --- | --- | --- | --- | --- | --- | --- | --- |
|  |  | GBS |  | GBM |  |  |  |  |  |
| Param. | Exact | Estimate | RMSE | Estimate | RMSE | Param. | Exact | Estimate | RMSE |
| LHx | 0.0302 | 0.0322 | 0.1250 | 0.0316 | 0.1774 | $\beta_{LHx}$ | 0.0302 | 0.0300 | 0.0071 |
| RAI | 0.0111 | 0.0110 | 0.0684 | 0.0102 | 0.1689 | $\beta_{RAI}$ | 0.0111 | 0.0111 | 0.0071 |
| KFk | 12.182 | 12.219 | 0.0270 | 11.579 | 0.0924 | $\beta_{KFk}$ | 12.182 | 12.121 | 0.0071 |
| KHk | 20.085 | 20.264 | 0.0941 | 20.594 | 0.0876 | $\beta_{KHk}$ | 20.085 | 19.985 | 0.0071 |
| OCp | 665.14 | 735.00 | 0.2535 | 614.85 | 0.1225 | $\beta_{OCp}$ | 665.14 | 661.81 | 0.0071 |
| RSS | 33.115 | 34.212 | 0.2004 | 30.249 | 0.1654 | $\beta_{RSS}$ | 33.115 | 32.950 | 0.0071 |
| TAS | 20.085 | 25.948 | 0.4684 | 20.244 | 0.1661 | $\beta_{TAS}$ | 20.085 | 19.985 | 0.0071 |
| TPF | 12.182 | 14.310 | 1.2457 | 11.945 | 0.1484 | $\beta_{TPF}$ | 12.182 | 12.121 | 0.0071 |
| RSO | 1.0101 | 0.9067 | 0.5032 | 0.8604 | 0.3126 | $\beta_{RSO}$ | 1.0101 | 1.0092 | 0.0011 |
| $\sigma^2$ | 0.05 | 0.0466 | 0.0034 | - | - | $\Gamma_{LHx}$ | 0.0030 | 0.0030 | 0.0002 |
| $\sigma_1^2$ | 0.05 | - | - | 0.0979 | 0.1383 | $\Gamma_{RAI}$ | 0.0011 | 0.0011 | 0.0016 |
| $\sigma_2^2$ | 0.05 | - | - | 0.1229 | 0.1726 | $\Gamma_{KFk}$ | 1.2182 | 1.2288 | 0.0012 |
| $\sigma_3^2$ | 0.05 | - | - | 0.1054 | 0.1488 | $\Gamma_{KHk}$ | 2.0086 | 2.0158 | 0.0005 |
| $\sigma_4^2$ | 0.05 | - | - | 0.1015 | 0.1434 | $\Gamma_{OCp}$ | 66.514 | 64.461 | 0.0044 |
| | | | | | | $\Gamma_{RSS}$ | 3.3115 | 3.3064 | 0.0002 |
| | | | | | | $\Gamma_{TAS}$ | 2.0086 | 2.0156 | 0.0005 |
| | | | | | | $\Gamma_{TPF}$ | 1.2182 | 1.2287 | 0.0012 |
| | | | | | | $\Gamma_{RSO}$ | 0.1010 | 0.1011 | 0.0001 |
| | | | | | | $\sigma^2$ | 0.0500 | 0.0697 | 0.0640 |

Both GB strategies resulted in larger RMSE values compared to PB. A large variability emerged among the RMSE values for individual parameters, with values ranging from 0.003 to 1.2. When compared to the estimations obtained by means of the single objective formulation (GBS/GA), the GBM/NSGA-II was generally less accurate, yielding larger or equivalent RMSE values. In particular, smaller RMSEs were obtained for parameters *RAI KFk, RSS* when GBS/GA was used, whereas RMSE for parameters *OCp, RSS, RSO* and *KHk* were comparable among the two GB formulations. One parameter, *TPF*, however, stood out as poorly predicted by the GBS/GA approach. It is interesting to notice that *TPF* corresponded to a very low-sensitive parameter of the GBS/GA objective function and this could explain that the algorithm wandered around a wide range of values without any impact on the prediction quality (see Supplementary figure S5). On the other hand, this parameter ranked third for the objective function of fructose and is involved in the interactions for the remaining sugars (Supplementary figure S4). Therefore, the GBM/NSGA-II better accounted for these multiple involvements and thus this could partly explain the difference between the GBS and GBM results in favour of the last one.

PB approach provided a reliable estimation of distribution parameters *θ*_*PB*_ as proved by the low RMSE values ranging from 1*e*^−4^ to 6*e*^−2^ (see Table 3). Good performances for the estimation of *θ*_*PB*_ also extended to the estimation of individual genotypic parameters. Figure 5 shows a comparison between estimated and true individual parameter values obtained for the three considered algorithms (GBS/GA, GBM/NSGA-II, PB/SAEM), out of the 100 simulated datasets. Results clearly point to a greater accurateness of the PB approach, with expected errors below 5% for all parameters. Among the GB approaches, the GBS formulation gives satisfactory results (expected error less than 10%) for most parameters but completely fails the estimation of parameters *TAS, TPF* and *RSO*, consistently with their high RMSE value. Results obtained with the GBM formulation are more homogeneous but of lower quality, with expected error close to 20%, for most parameters.

**Figure 5:**
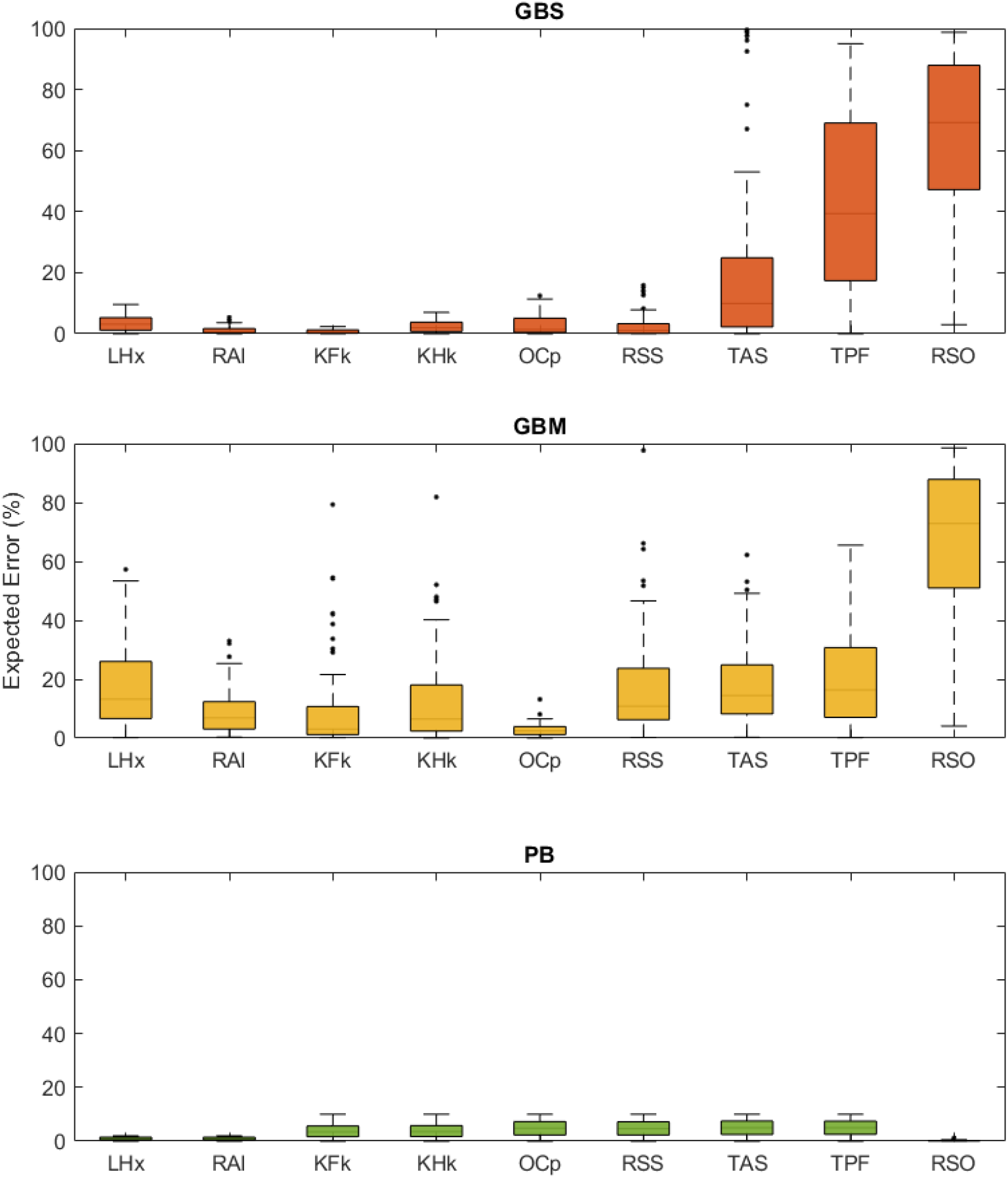
Expected error (%) on the individual estimates of GBS/GA, GBM/NSGA-II and PB/SAEM approach over 100 simulated datasets.

### 6.3 Population-based approach provides robust estimates on real data

Results on simulated data were confirmed by estimations obtained over the peach progeny. When comparing results obtained over different repetitions of the calibration algorithm, GB strategies suffered from a substantial lack of reproducibility.

Figure 6 shows the relative standard deviation of *KHk* estimates obtained for the 106 genotypes, with the three considered approaches. The distribution is a lot wider for GB approaches compared to the PB strategy. The robustness of GBM estimations, in particular, strongly depended on the considered genotype, with standard deviation values ranging from zero to 70%. When comparing individual parameters, large variations in the estimated values were observed for parameters *TAS, TPF*, *OCp* and *RSS* (see Supplementary figure S3), although these parameters do not always correspond to high expected RMSE values (see Table 3). In agreement with the simulation study, the PB approach proved much more stable across repetitions, independently of the genotype (Fig 6). Table 4 reports the confidence intervals of the vector of parameters *θ*_*PB*_, characterizing the inter-individual distribution. For all estimated parameters, the confidence intervals are remarkably narrow, confirming the ability of PB approach to robustly estimate both mean and individual parameter values of the population.

**Table 4:** PB approach: results of the estimation of mean population parameters *θ*_*PB*_ over the real peach progeny. Comparison to parameter mean and variance obtained from individual GB estimates.

|  | PB |  |  | GBS | GBM |
| --- | --- | --- | --- | --- | --- |
| Parameter | Estimate | SD | 95% CI | Estimate | Estimate |
| $\beta_{LHx}$ | 0.4914 | 0.0093 | [0.4747; 0.5108] | 0.45 | 0.42 |
| $\beta_{RAI}$ | 0.1075 | 0.0005 | [0.1065; 0.1086] | 0.028 | 0.028 |
| $\beta_{KFk}$ | 34.8449 | 0.8078 | [33.0419; 36.2398] | 34.7 | 45.1 |
| $\beta_{KHk}$ | 206.3626 | 2.9419 | [200.8797; 212.3122] | 236.0 | 251.9 |
| $\beta_{OCp}$ | 774.9177 | 4.8343 | [765.3165; 784.1586] | 943.9 | 1001 |
| $\beta_{RSS}$ | 406.2915 | 4.1089 | [398.8225; 414.7968] | 449.4 | 326.0 |
| $\beta_{TAS}$ | 147.5079 | 1.4867 | [144.3786; 150.1947] | 184.0 | 136.7 |
| $\beta_{TPF}$ | 51.0577 | 0.7856 | [49.2564; 52.4168] | 25.7 | 70.38 |
| $\beta_{RSO}$ | 2.6561 | 0.0975 | [2.4496; 2.8277] | 2.21 | 2.66 |
| $\Gamma_{LHx}$ | 1.9430 | 0.0355 | [1.8704; 2.0099] | 161.7 | 2.07 |
| $\Gamma_{RAI}$ | 2.7355 | 0.0208 | [2.6967; 2.7764] | 72.2 | 2.62 |
| $\Gamma_{KFk}$ | 2.0338 | 0.0260 | [1.9884; 2.0921] | 2.82 | 6.51 |
| $\Gamma_{KHk}$ | 1.9708 | 0.0268 | [1.9174; 2.0210] | 1.05 | 1.04 |
| $\Gamma_{OCp}$ | 1.2225 | 0.0019 | [1.2189; 1.2266] | 1.04 | 1.01 |
| $\Gamma_{RSS}$ | 2.0764 | 0.0239 | [2.0278; 2.1224] | 33.1 | 1.76 |
| $\Gamma_{TAS}$ | 2.2615 | 0.0258 | [2.2150; 2.3154] | 3.82 | 1.76 |
| $\Gamma_{TPF}$ | 2.4144 | 0.0360 | [2.3521; 2.4936] | 9.93 | 1.46 |
| $\Gamma_{RSO}$ | 2.7073 | 0.0323 | [2.6427; 2.7705] | 247.6 | 1.36 |
| $\sigma^2$ | 0.9824 | 0.0341 | [0.9199; 1.0484] | - | - |

**Figure 6:**
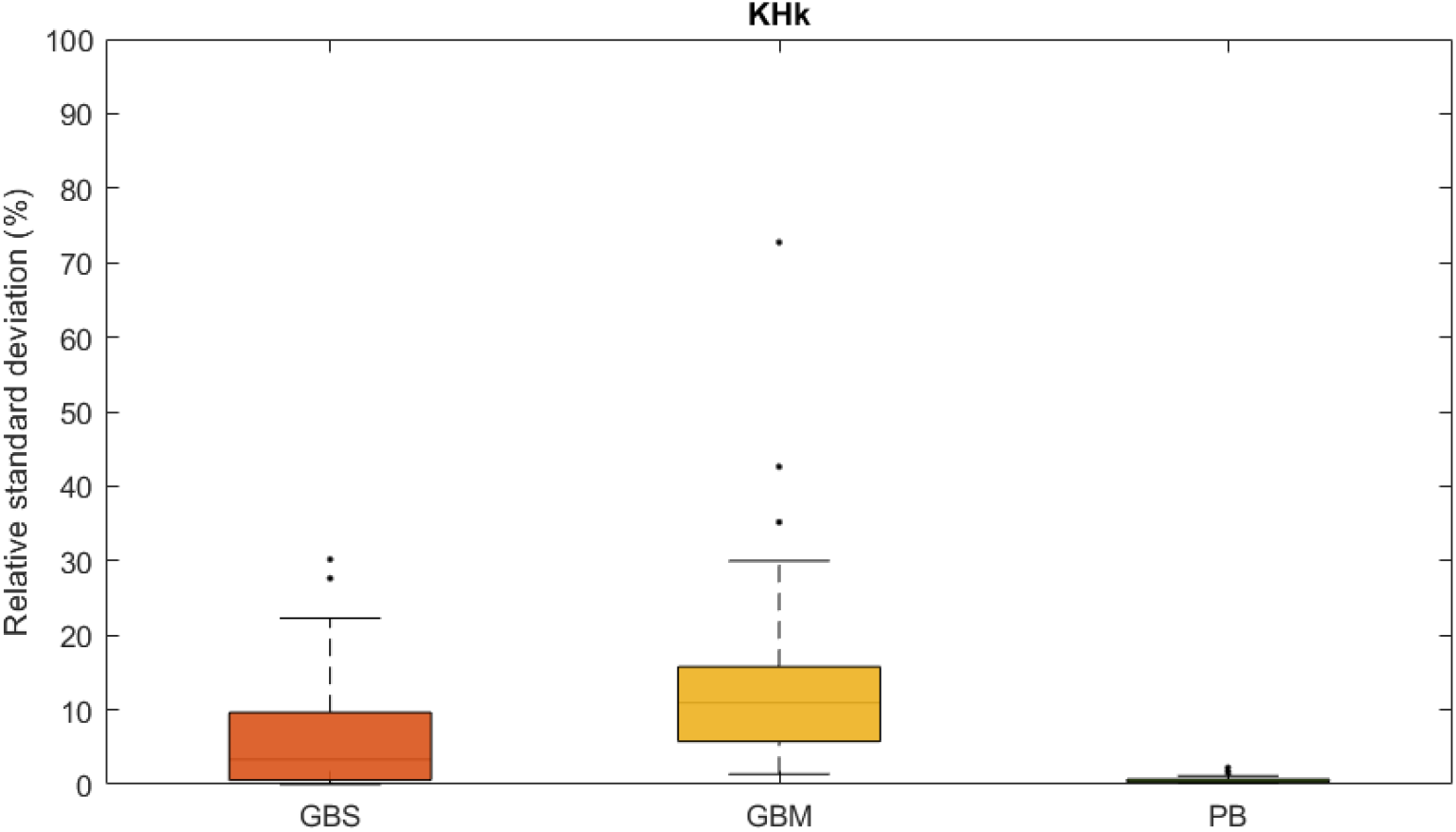
Estimation variability over repetitions of the calibration algorithm. Relative standard deviation of individual estimates of the parameter KHk over the 106 genotypes using GBS/GA, GBM/NSGA-II and PB/MCMC-SAEM.

### 6.4 Candidate role of Kfk in low-fructose phenotype is confirmed over the whole peach population

Quantifying the inter-individual variability of model parameters is an important objective of genetic studies. Table 4 compares the population parameters computed from individual GB estimates to the *θ* values by PB approach. Results show GB and PB approaches overall agree on the estimation of mean parameters (*β*), at least in their magnitude, providing a good overview of the timescales of the system. Conversely, the estimation of parameter variations around the mean value turns out to be more sensitive to the calibration method. Although estimations with PB and GBM approaches share similar orders of magnitude, quantitative difference are observed in the estimated variance for several parameters. The GBM strategy in particular usually results into a lower parameter variability with respect to the PB approach. The worst results, however, are obtained with the GBS approach. Single-objective calibration largely over-estimate the expected variance for parameters *LHx, RAI, RSS, TPF, RSO*, as the result of a very right-skewed distribution (Figure S6). This is probably due to a handful of genotypes for which the estimation process is unstable.

Despite the aforementioned discrepancies, both GB and PB methods were able to capture specific features of the peach population under study. For all three methods, the distribution of *KFk* was markly bimodal, suggesting the presence of two sub-populations, with contrasted affinity values. This result is in agreement with the prediction of the model (Desnoues et al., 2018) according to which the fructokinase affinity may be responsible for the appearance of two distinct fructose phenotypes, as observed in the peach population. In order to further investigate the relationships between the parameter values and the resulting fructose concentration, we systematically compared the estimations obtained over the two phenotypic populations, with the three calibration methods (see Figure 7). Results supported a significative and robust difference in the estimated values of *KFk* for the two phenotypic groups, corroborating the involvement of the fructokinase affinity in the specification of the fructose content of the fruit. Other parameters emerged as different between the two phenotypic groups when considering the results from the GB calibration methods (see Figure S7), but these results were not supported when using the PB approach. This was the case for example for *KHk* and *TPF*, which were also pointed out as putative mechanisms in the original modeling work by Desnoues et al. (2018), using a GBS approach. These results underline the importance of the choice of the calibration method which may impact the biological interpretation of the results.

**Figure 7:**
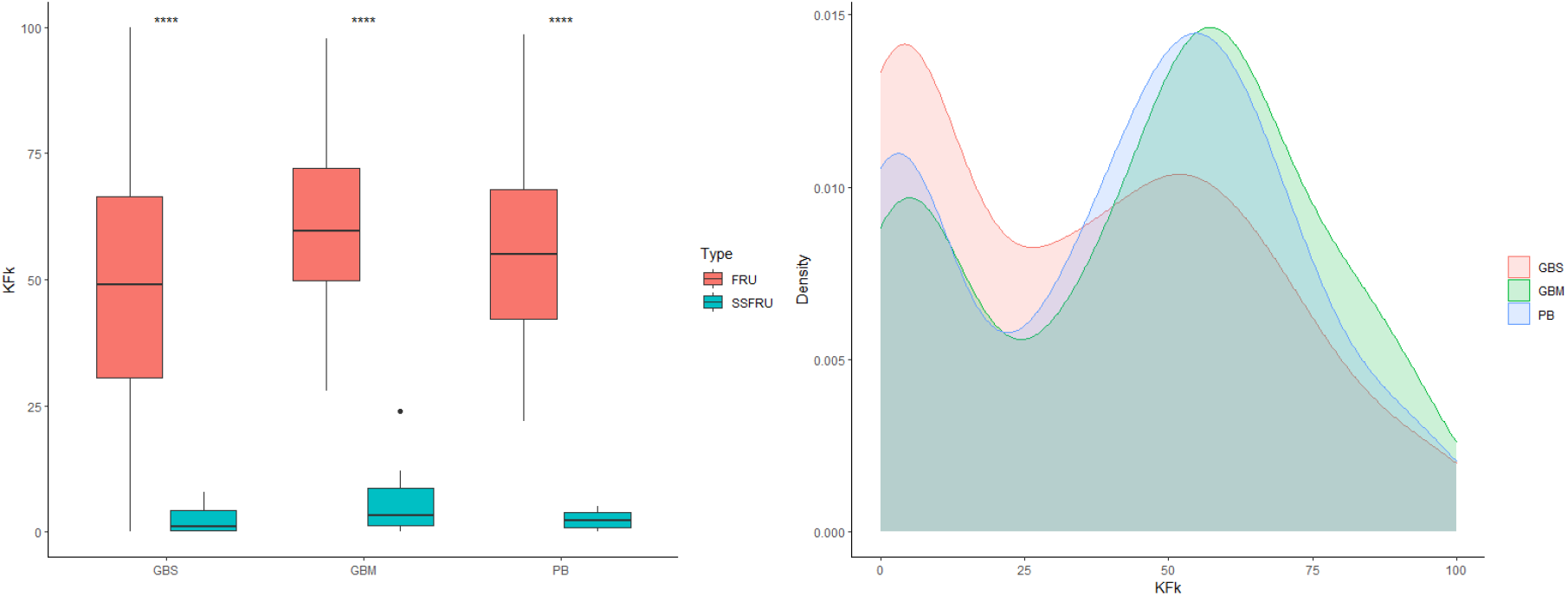
Difference in the estimated model parameter (KFk) between standard (‘FRU’) and low fructose phenotypes (‘SSFRU’) and its distribution over 106 genotypes obtained with the three considered methods: GBS, GBM and PB approach.

## 7 Discussion

In the context of agronomy, models are increasingly used to describe plant growth and metabolism and test their performances in different conditions. Being able to account for intra-specific variability is a key challenge in the field as it can provide useful information for varietal selection and plant breeding. In this paper, a mathematical model of fruit sugar metabolism (Kanso et al., 2020) was calibrated over a population of 106 peach varieties using two contrasted strategies that strongly differ in the way data are interpreted and analysed: an individual, genotype-based (GB) approach and a population-based (PB) approach. The former was further decomposed into a single-objective (GBS) and a multi-objective (GBM) formulation, in order to account for possible trade-off among phenotypic traits.

When applied to the model proposed by Kanso et al. (2020), all approaches led to a satisfactory agreement between predictions and data. In term of phenotypic predictions, the GBS formulation generally provided better results, especially concerning the dynamics of sucrose and glucose, the prevailing sugars in peach fruit, closely followed by PB approach. However, when looking at the estimation of the underlying parameter values, the GB strategy suffered from a lack of reproducibility, with several parameters sets giving an equivalent agreement with data. This is a well-known issue of collective fit in system biology: in spite of a good predictive power, parameter estimation may show large uncertainty, in correspondence to sloppy direction of the parameter space (Gutenkunst et al., 2007). Tests over a simulated dataset showed that, in the case of GBS, sloppiness mainly concerned three (out of nine) model parameters, *TAS, TPF, RSO*, for which estimates were both inaccurate (see Table 3, Fig.5) and unstable across repetitions (see Fig.S3). The use of a multi-objectives (GBM) formulation did not really improve results, resulting into an intermediate, but still large variability of estimates. Indeed, although the contribution of individual sugars to the objective function was better represented, the inclusion of 4 additional parameters (*σ*_*i*_) reduced the number of data-points available for the parameter estimate. Moreover, it could be worth mentioning here that NSGA-II is known to be particularly effective for optimisation problems involving two or three objective functions but that its performances decrease as the dimension increases (Wang et al., 2013).

Better results were undoubtedly obtained with the population-based approach. The advantage of this approach is that it takes into account the observation of the whole progeny and cast them in the same general framework (common probability distribution). The presence of multiple genotypes with a similar sugar concentration indeed helped to constraint the estimation process, resulting in a high stability and accuracy of individual estimates, on both the simulated (Figure 5) and the experimental datasets (Figure 6 and Table 4).

In addition to individual estimates, PB approaches naturally yield information on parameter distribution across genotypes. In particular, it allows for a direct analysis of genetically-relevant correlations among parameters, via the estimation of the covariance matrix. To this aim, different covariance structure can be tested, reflecting available information or alternative biological hypotheses. For the peach progeny used in this studied, a diagonal covariance matrix, corresponding to the parameter independence, proved the best choice according to the AIC criterion (Burnham and Anderson, 2002), compared to a full-rank matrix (full covariance matrix *AIC* = 2396.1 vs diagonal covariance *AIC* = 2299.7). This suggests that the genetic control of the biochemical reactions involved in peach sugar metabolism is split over independent regulatory modules (Desnoues et al., 2016; Carreno-Quintero et al., 2013).

When comparing PB results to parameter distribution inferred from individual GB-based estimates, results showed that both strategies overall agreed on the estimation of mean parameter values (Table 4) but the estimation of parameter variance was strongly affected by the presence of outliers in the case of GB approaches. In spite of this limitation, both GB and PB approaches and were able to capture specific features of our peach population. The estimation of parameter *KFk*, corresponding to the affinity of fructokinase, revealed a bimodal distribution, i.e. the observed variability corresponded to the presence of two sub-populations, with different mean values. This result agrees remarkably well with the existence of two distinct phenotypes in the population, with contrasted fructose contents, and with previous analysis (Desnoues et al., 2018) suggesting *KFk* as the main determinant of such a phenotypic trait.

In light of our results, the PB approach appears a suitable choice for genetic studies, as long as a few conditions are satisfied. First, the advantages of PB strategy with respect to GB approaches may depend on the population size. In Baey et al. (2018), parameter estimation over a population of 34 genotypes gave rise to instabilities, due to convergence issues, and proved sensitive to the parameters setting of the algorithm. In our case, the large population size ensured a good and stable estimation of parameter values, independently of the starting conditions and the algorithm setting. For the tuning parameters there is no general recommendation in the literature, and in practice one may have to run the algorithm several times to identify reasonable values for these parameters. For populations of small size, GB approaches offer a good alternative in terms of phenotypic prediction but the algorithm has to repeated several times in order to check the stability of the estimates. In case of problems, efforts should be directed in improving the number of datapoints, in order to further constraint the exploration of parameter space. Last but not least, the fundamental hypothesis made in this paper for the PB approach, namely that individual genetic parameters are realization of a Gaussian distribution, might not be valid when multi-specific or natural (inbred) populations are considered. In this case, extension of the approach can be considered, with non-Gaussian distribution for the individual parameters, accounting for the presence of sub-populations. However, parameter estimation is more involved in these cases, and other numerical methods may be necessary, such as Bayesian approaches for example.

At term, a reliable estimation of model parameters will offer the possibility to study the genetic control of these parameters, looking for genomic regions (QTL) of interest (Cooper et al., 2009; Yin et al., 2016). This will open new opportunities for the analysis of genotype x environment interaction and virtual breeding *i*.*e*. the design of new genotypes that better meet the challenges of modern agriculture (e.g. increased production, new cultivation techniques, future climates).

## Acknowledgments

We would like to thank V. Signoret for her help in maintaining the inter-specific peach progeny. We are grateful to the IE-EMMAH UMR1114 and IE-GAFL UR1052 teams for taking care of the experimental orchard.

